# Targeting the fronto-parietal network using multifocal personalized transcranial alternating current stimulation to enhance motor sequence learning in healthy older adults

**DOI:** 10.1101/2022.02.16.480660

**Authors:** L.R. Draaisma, M.J. Wessel, M. Moyne, T. Morishita, F.C. Hummel

**Author notes:** Correspondence: Prof. Dr. Friedhelm C. Hummel, Defitech Chair of Clinical Neuroengineering, Center for Neuroprosthetics (CNP) and Brain Mind Institute (BMI), Swiss Federal Institute of Technology (EPFL), 9 Chemin des Mines, 1202 Geneva. Declaration of interest: none.

## Abstract

**Background:** Healthy older adults show a decrease in motor learning capacity as well as in working memory (WM) performance. WM has been suggested to be involved in motor learning processes, such as sequence learning. Correlational evidence has shown the involvement of the fronto-parietal network (FPN), a network underlying WM processes, in motor sequence learning. However, causal evidence is currently lacking. Non-invasive brain stimulation (NIBS) studies have focused so far predominantly on motor related areas to enhance motor sequence learning while areas associated with more cognitive aspects of motor learning have not yet been addressed.

**Hypothesis:** In this study, we aim to provide causal evidence for the involvement of WM processes and the underlying FPN in successful motor sequence learning by using a theta transcranial alternating current stimulation (tACS) paradigm targeting the FPN during motor sequence learning.

**Methods:** In a cohort of 20 healthy older adults, we applied bifocal tACS in the theta range to the FPN during a sequence learning task. With the use of a double-blind, cross-over design, we tested the efficacy of active compared with sham stimulation. Two versions of the motor task were used: one with high and one with low WM load, to explore the efficacy of stimulation on tasks differing in WM demand. Additionally, the effects of stimulation on WM performance were addressed using an N-back task. The tACS frequency was personalized by means of EEG measuring the individual theta peak frequency during the N-back task.

**Results:** The application of personalized theta tACS to the FPN improved performance on the motor sequence learning task with high WM load (*p* <.001), but not with low WM load. Active stimulation significantly improved both speed (*p* <.001), and accuracy (*p* =.03) during the task with high WM load. In addition, the stimulation paradigm improved performance on the N-back task for the 2-back task (*p* = .013), but not for 1-back and 3-back.

**Conclusion:** Motor sequence learning can be enhanced with the use of personalized bifocal theta tACS to the FPN when WM load is high. This indicates that the efficacy of this stimulation paradigm is dependent on the cognitive demand during the learning task and provides further causal evidence for the critical involvement of WM processes and the FPN in motor sequence learning in healthy older adults. These findings open new exciting possibilities to counteract the age-related decline in motor learning capacity and WM performance.

## Introduction

The ability to acquire new motor skills is important in daily life. Motor learning is a practice-dependent process in which consequence movements are performed quicker and more accurately [1]. A vast amount of research has contributed to an increased understanding of the neural substrates and underlying mechanisms involved in the acquisition, consolidation and retention of new motor skills. Neuroscientific studies have focused predominantly on the motor network and the pivotal role of the primary motor cortex (M1) [2–4]. This is especially the case for non-invasive brain stimulation (NIBS) studies that attempt to improve motor learning by combining the practice of a challenging motor task with a stimulation paradigm [5–8]. However, studies have suggested that challenging motor tasks, such as motor sequence learning (MSL), do not rely exclusively on motor related processes, but also on cognitive processes, such as working memory (WM) processes [4,9,10]. Surprisingly, WM related brain areas have not been a target for NIBS paradigms intended to study MSL.

MSL is a process where independent movements are associated, eventually resulting into a multi-element sequence that can be performed quickly and accurately [4,11]. Studies have shown the involvement of WM in MSL [12–14]. WM refers to the ability to temporarily store and manipulate information in the mind [15]. Inter-individual variability in WM consist of the number of items that can be held and worked with [4]. This is important for MSL especially during the process of grouping elements of the sequence together in “chunks”. This chunking process results into quicker execution of the movements [16–18]. Many studies have shown that healthy older adults show a decline in the ability to learn motor sequences [19,20]. Moreover, aging decreases cognitive functions including WM [21]. Therefore, an interaction among age, WM capacity and MSL has been recently suggested [19], though causal evidence in favour of this suggestion remains limited.

A promising neurotechnology to provide causal evidence is the use of NIBS such as transcranial alternating current stimulation (tACS) [22–25]. This technique allows to exogenously interfere with ongoing oscillatory activity and to target specific networks, such as the fronto-parietal network (FPN), to enhance or decrease specifically respective cognitive functions, such as WM processes [26,27]. The FPN, a network related to WM, has shown to be activated during motor sequence tasks [16,28–31]. Cognitive processes rely on coordinated interactions within and among brain networks, implemented in the brain by oscillatory activity [32,33]. For example, efficiency is increased by oscillatory synchronization of neuronal firing, which creates ensembles of neurons that carry out specific computational functions [33,34]. The main working mechanism of tACS is to entrain with or synchronize neuronal networks [22,35]. The stimulation frequency is adjusted to match the endogenous oscillatory frequency and its brain state. More specifically, tACS allows to exogenously interact with ongoing oscillations, which can result in enhanced coherence within networks with the respective behavioural impact [25,36,37]. Neuronal oscillations in the theta range (4-8 Hz) are engaged in WM tasks, with an increase in theta power during increased WM load [38–40]. Polania and colleagues and Violante and colleagues have shown a causal relationship between the synchronization of theta oscillations with a relative 0° phase difference in the FPN and the improvement of WM performance [26,27]. However, knowledge about the effects of tACS induced synchronization of theta oscillations in the FPN and MSL is lacking.

In this study, we aimed to determine a causal relationship between WM and MSL in healthy older adults. To do this, personalized theta tACS was applied to the right dorsolateral prefrontal cortex (DLPFC) and the posterior parietal cortex (PPC) intended to improve MSL by means of the sequential finger tapping task (SFTT) [3]. To evaluate the importance of WM during MSL and how this is affected by the FPN stimulation, two versions of the SFTT were used. The versions differed in terms of low vs. high WM load. WM load was kept low by explicitly showing the sequence on a screen during the task [41]. In the high WM load version, the sequence had to be memorized prior to the task and was not shown during the task. This online maintenance of the sequence while performing the movements relies relevantly on WM processes [42]. With this study, we introduce the FPN as an additional stimulation target location for motor learning enhancement and shine a light on the importance of taking cognitive processes into account during MSL paradigms.

## Methods

### Participants

In this study, we recruited N = 21 healthy, older, right-handed participants (N = 11 female, mean age ± sd: 69.6 ± 4.4, mean laterality quotient Edinburgh handedness inventory 85.03 ± 17.3 [43]. The data of N = 20 participants were considered due to a drop-out of one participant caused by an unrelated change in physical health. Inclusion criteria were: ≥ 60 years, right-handed and absence of contraindications for transcranial electrical stimulation (tES). Exclusion criteria were: neuropsychiatric diseases, history of seizures, medication that potentially interacts with tES, musculoskeletal dysfunction that impairs finger movements, professional musician, intake of narcotic drugs and the request of not being informed in case of incidental findings. All participants have signed an informed consent. The study was performed in accordance to the declaration of Helsinki [44]. Ethical approval was obtained from the cantonal ethics committee Vaud, Switzerland (project number: 2017-00765).

### Experimental design

The design of this study was double blind, sham-controlled, cross-over. It consisted of two sessions before cross-over and two sessions after cross-over. During the session on day 1, the participants were informed, screened and asked to fill in three different questionnaires (tES safety questionnaire, Edinburgh Handedness Inventory (EHI), Center for Epidemiological Studies Depression Scale (CES-D)) [43,45]. Afterwards the participants performed an N-back test with EEG acquisition for peak frequency analysis. Following the EEG measurement, the participants did the motor training and the cognitive training with concurrent tACS. The next day the participants performed the motor task with tACS. Between the before and after cross-over sessions there was a minimum time period of two weeks. The same tasks, with different sequences, were repeated after cross-over, excluding the questionnaires. Please see figure 1 A for the timeline of study design.

**Figure 1.**
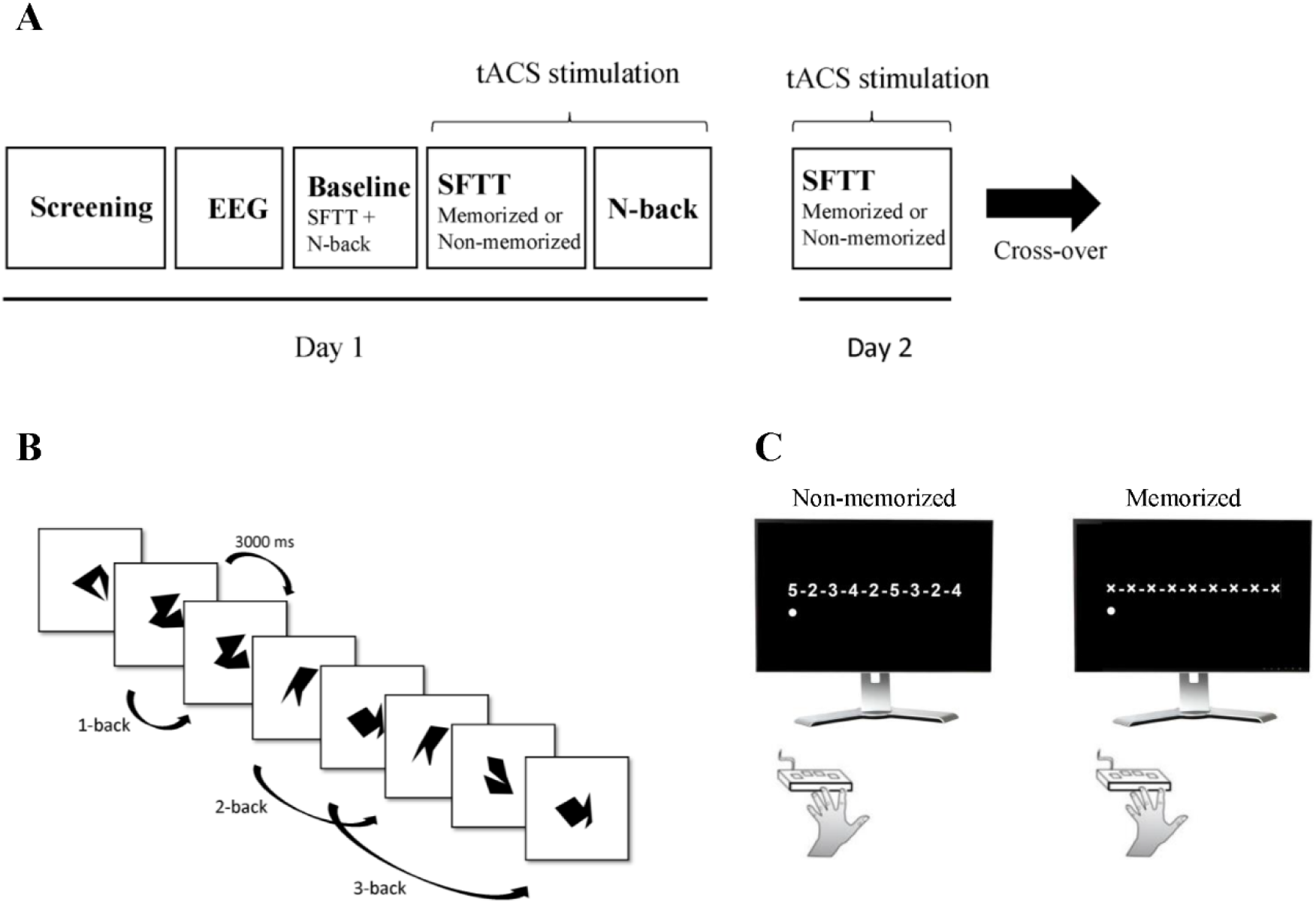
Experimental design. **A)** timeline of study design. **B)** Example of N-back test with the three difficulty levels shown. **C)** Example of the non-memorized and the memorized version of the SFTT. Please note that with each key press advancing point indicates in both conditions just the position within the sequence

### Cognitive task

Participants were asked to perform the N-back task. The task was implemented in Matlab (The MathWorks Inc.,Natick, Massachusetts, USA) and was based on the single N-back task used by Jaeggi and colleagues [46]. The script was adapted from Quent, A.J. [47] in terms of language (French & English), length and difficulty level. The task consisted of a sequence of visual stimuli that were shown on a computer screen. The participants had to respond by clicking the right “Control” button on a computer keyboard when the stimulus was the same as the stimulus presented N positions back. Participants should not respond when a different stimulus was presented. The visual stimuli consisted of 10 random shapes, eight 8-points shapes (number 14, 15, 17, 18, 20, 22, 23, and 27) and two 12-points shapes (number 20 and 24) taken from Vanderplas and Garvin [48]. The stimuli were presented for 500 ms each with a 2500 ms interstimulus interval. The participants were required to respond within the response window that starts at the onset of the stimulus until the end of the interstimulus interval (3000 ms). The task consisted of 1 until 3-back levels, in that order. The task was divided into a baseline and training session, with the baseline session consisting of 1 block per n-back level (3 blocks in total) and the training session of 3 blocks per level (9 blocks in total). Every block consisted of 20 + n trials, with 6 targets and 14 + n non-targets. The reaction times, hits, misses, false alarms and correct rejections were measured. Please see figure 1 B for a schematic illustration of the task. We had to exclude N = 9 before cross-over N-back task data sets due to an error in the response recording. A total of N =31 N-back data sets were considered.

### Motor learning task

Participants executed two different versions of the sequential finger tapping task (SFTT) based on the SFTT task used in earlier studies [3,49]. They were asked to perform a 9-item sequence with their non-dominant left hand. They were orally instructed to continuously tap the same sequence as fast and as accurately as possible on a four-button keyboard (Current Designs, Philadelphia, PA, USA). The sequences consisted of 4 digits from 2 to 5, which corresponded to the four fingers from index (2) to the little finger (5) of the left hand. Different sequences were used for the baseline and the training measurements. All sequences were matched in complexity verified with the Kolmogorov complexity test [50]. The baseline measurement consisted of one block of 90 seconds, the training measurement consisted of seven blocks of 90 seconds, with 90 seconds breaks after every block which lasted 20 minutes in total. The task was implemented in Presentation software (Neurobehavioral Systems, Berkeley, CA, USA). Participants performed a low and a high WM load SFTT version. The low WM load version displayed the sequence on the screen asking participants to execute the sequence without prior familiarization. A cursor underneath the displayed sequence moved in response to every finger tap to identify the target digit. This version is referred to as the “non-memorized” version. For the high WM load version, participants had to memorize the sequence before the task started. They received the sequence on a paper and were asked to learn the sequence by heart, without practicing it on the button keyboard. With the use of a distractor task, during which they had to spell random words in a reversed order sufficient memorization of the sequence was verified. More precisely, participants had to spell backwards 3 words in a row, and recall the sequence out loud afterwards. After 3 times correct, the sequence was deemed sufficiently learned [51]. During the memorized version of the task, the participants could not see the sequence. Displayed on the screen was a sequence of 9 “X’s” with the moving cursor underneath to identify the target digit. Participants performed both versions divided over day 1 and day 2 in a randomized order. This order of versions was reversed after cross-over, see figure 1 C.

### Transcranial alternating current stimulation

Multifocal tACS was applied to the right FPN using two neuroConn DC plus stimulators to enable bifocal stimulation (neuroConn GmbH, Ilmenau, Germany). Participants received both real (30 min) or sham (30 seconds) stimulation in randomized order, before or after cross-over [52]. The stimulation protocol consisted of the following parameters: in-phase (0° phase lag), intensity 2 mA (peak-to-peak), fade-in/fade-out interval of 8 seconds. The stimulation frequency was adjusted to the personal theta peak frequency, which was recorded during an EEG recording while performing a pre-baseline N-back test of 1 block per level. Rubber concentric electrodes were used: centre electrode size diameter: ca. 20 mm, area: ca. 3 cm^2^ and ring electrode size diameter: out 100 mm/ in 70mm, area: ca. 40 cm^2^. Electrode location was defined with the use of a standard 64 channel, EEG actiCAP with 10/20 system (Brain Products GmbH), targeting F4 and P4. The paste used for conductivity with adequately low impedance was SAC2 electrode cream (Spes Medical Srl, Genova, Italy). This paste was adhesive which ensured stable electrode placements. The electrode placement and the electric field distribution were visualised with the use of standard template in SimNIBS (Version 3.2) [53]. The script to implement bifocal stimulation with ring electrodes was adapted from the open access Matlab script (© G. Saturnino, 2018). A template head model was used to simulate the electrode placement and electric field distribution. For the electrode placement and electric field distribution, please see figure 2 A & B.

**Figure 2.**
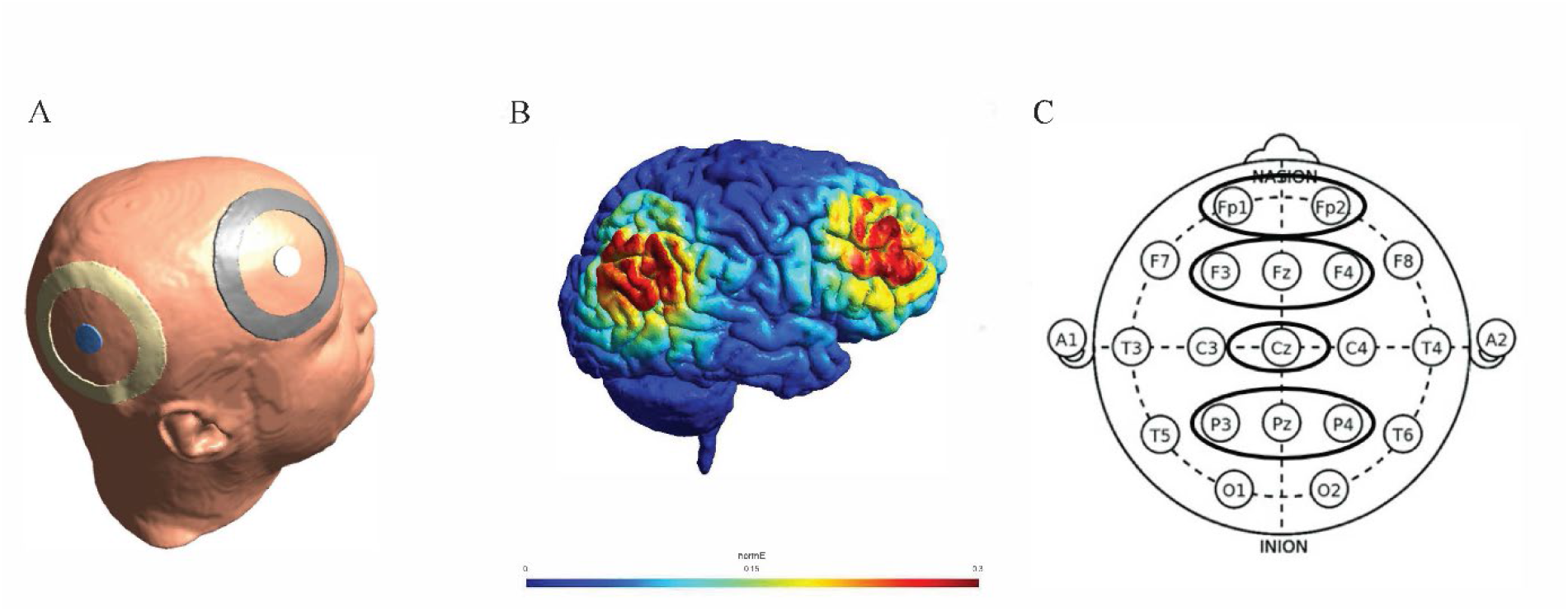
Bifocal tACS application and EEG recording. **A**) Bifocal electrode placement for tACS with concentric electrodes placed on F4 and P4. Image created with the use of SimNIBS software. Head is derived from a standard template provided. **B)** Simulation of the electric field distribution of tACS set-up created with the SimNIBS software. Label indicates strength of electric field (V/m). Brain is derived from a standard template provided in the program. Stimulation parameters are adjusted to the current study. **C)** EEG recording sites for the determination of the individual peak frequency during the working memory task.

### EEG

All EEG recordings were done in a shielded faraday cage. A customized electrode set-up with 9 electrodes was used, Frontal (Fp1, Fp2, F3, Fz, F4), parietal (Cz, P3, Pz, P4), please see figure 2 C. Using a 64-channel ANT Neuro EEG cap with eego™mylab software (ANT Neuro, Netherlands). EEG was recorded during the performance of the N-back task. With markers, the beginning and the end of every separate N-back level were defined. Recordings were done during 3 N-back blocks resulting in approximately 3 minutes of recording time. The peak frequency in the theta range (4 – 8 Hz) was calculated using a custom Matlab script (The MathWorks Inc., USA) adapted from the script used by Salamanca-Giron and colleagues [54] and made suitable for theta frequency analysis during N-back task performance.

### Data analyses

Normality of the data was visually checked with histograms and Q-Q plots of residual values and confirmed by verification of skewness ranging between 1 and -1 [55]. P-values of <.05 indicate statistical significance. Pre-processing of the behavioural data of the SFTT was done with an in-house script implemented in Matlab. Main output measures were: correct sequences, total completed sequences and correct sequences / completed sequences. Pre-processing of the individual N-back data was done with RStudio (version 1.4.1717, 2021)[56]. Individual data were combined in one main file using Microsoft Excel. All other analysis of the SFTT and the N-back data were done in Rstudio (version 1.4.1717). Data were analysed with the use of Linear mixed-effects models that were fitted with the “lmerTest” package. Output was type III anova table with p-values for F-tests [57]. Effect size was determined using partial eta squared with the “effectsize” package. Post-hoc analysis was done by pairwise comparisons, using the estimated marginal means and Tukey correction. Analysis of the tACS stimulation sensations and blinding responses were analysed with JASP (version 0.8.5.1) [58].

Responses to the real vs. sham stimulation estimations were analysed using a binomial test. The stimulation sensations were analysed using contingency tables with chi-squared analysis to control for differences between the real and sham stimulation conditions.

## Results

### Sequential finger tapping task

The two SFTT’s have been analysed separately as they both measure different types of motor learning (with high and low WM load). With the use of a linear mixed effects model the analysis of the amount of correct sequences of the **memorized** version of the SFTT showed a significant effect for blocks F(6, 247) = 18.57, *p* < .001, η_p_^2^ = 0.31 indicating a large effect size, as well as a significant effect for stimulation F(1, 247) = 18.83, *p* <.001, η_p_^2^ = 0.07 with a medium effect size, but no blocks x stimulation interaction F(6, 247) = 0.77, *p* = 0.59, η_p_^2^ = 0.02. The results of the **non-memorized** version show a significant effect for blocks F(6, 247) = 16.00, *p* <.001, η_p_^2^ = 0.28 (large effect), but no stimulation F(1, 247) = 0.46, *p* = .499, η_p_^2^ = 0.002 or interaction effect F(6, 247) = 0.36, *p* = .901, η_p_^2^ = 0.009. In an additional analysis the two conditions were combined to see whether there was a difference in the performance during block 1 in the sham condition. A pairwise t-test showed no significant difference between the memorized vs. the non-memorized condition t(19) = -0.34, *p* = .735, d = 0.08, indicating that participants showed a similar level of performance during block 1 for both conditions. After active stimulation only the memorized condition showed a trend for improvement compared with sham stimulation in block 7 t(19) = -2.07, *p* = .052, d = -0.46. Indicating that in both conditions, participants learned significantly but only in the memorized condition the tACS stimulation significantly improved performance, see figure 3.

**Figure 3.**
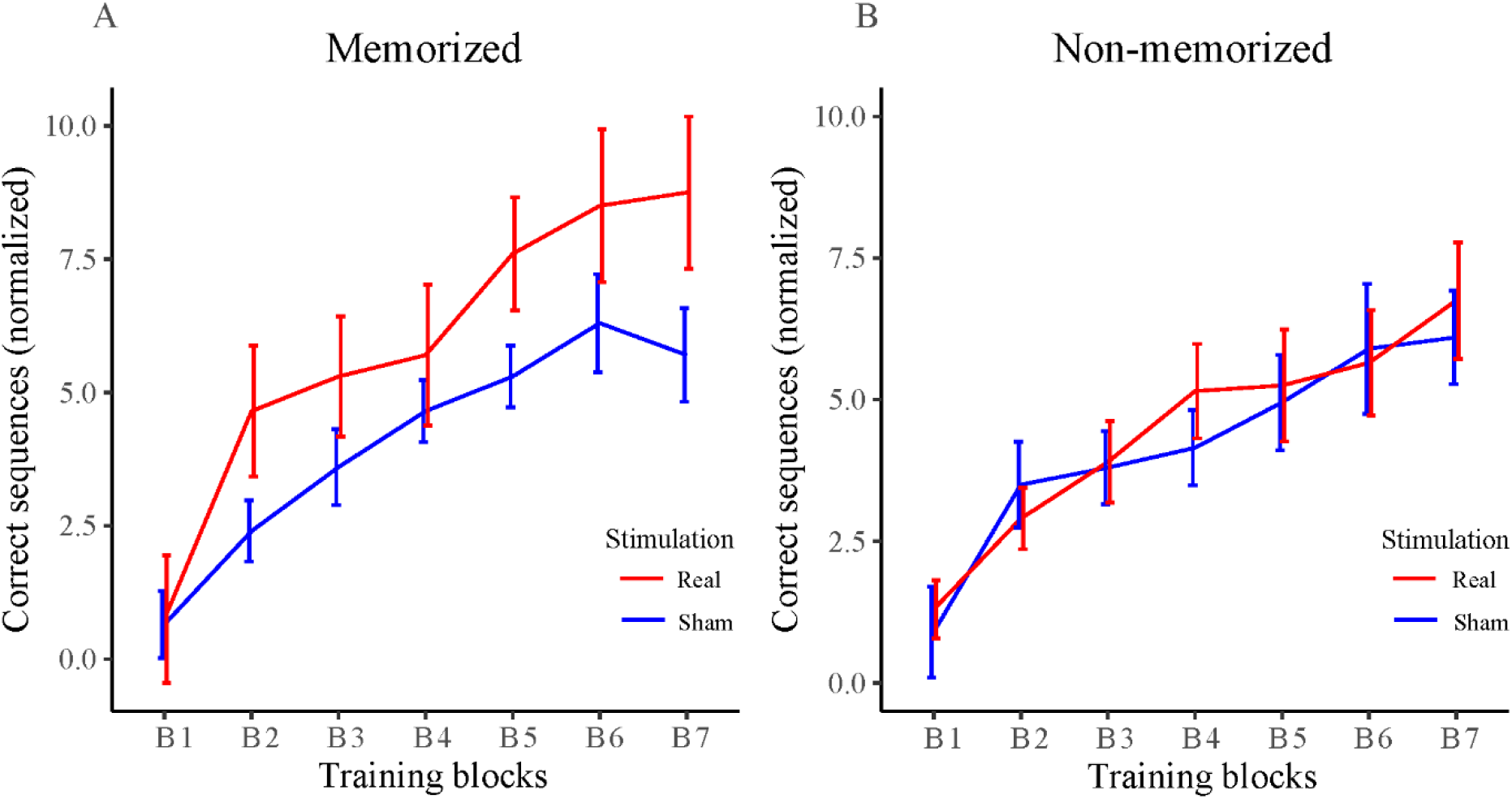
Plot of the correct sequences of the SFTT, results are normalized to baseline by subtraction. Error bars show standard error of the mean (SEM). On the left, the results of the memorized version. On the right, the results of the non-memorized version. Red lines depict the real stimulation and blue lines are the sham stimulation. Please note a significant stimulation effect with enhanced behavioural improvement in the memorized version (left graph).

#### Speed and accuracy

To further investigate the results of the **memorized** condition, the total amount of completed sequences were analysed as a measure of speed. The results showed a significant block effect F(6, 247) = 28.21, *p* < .001, η_p_^2^ = 0.41 (large effect) and a significant stimulation effect F(1, 247) = 15.92, *p* <.001, η_p_^2^ = 0.06 (small effect), but no interaction effect F(6, 247) = 0.23, *p* = 0.968, η_p_^2^ = 0.006, see figure 4A.

**Figure 4.**
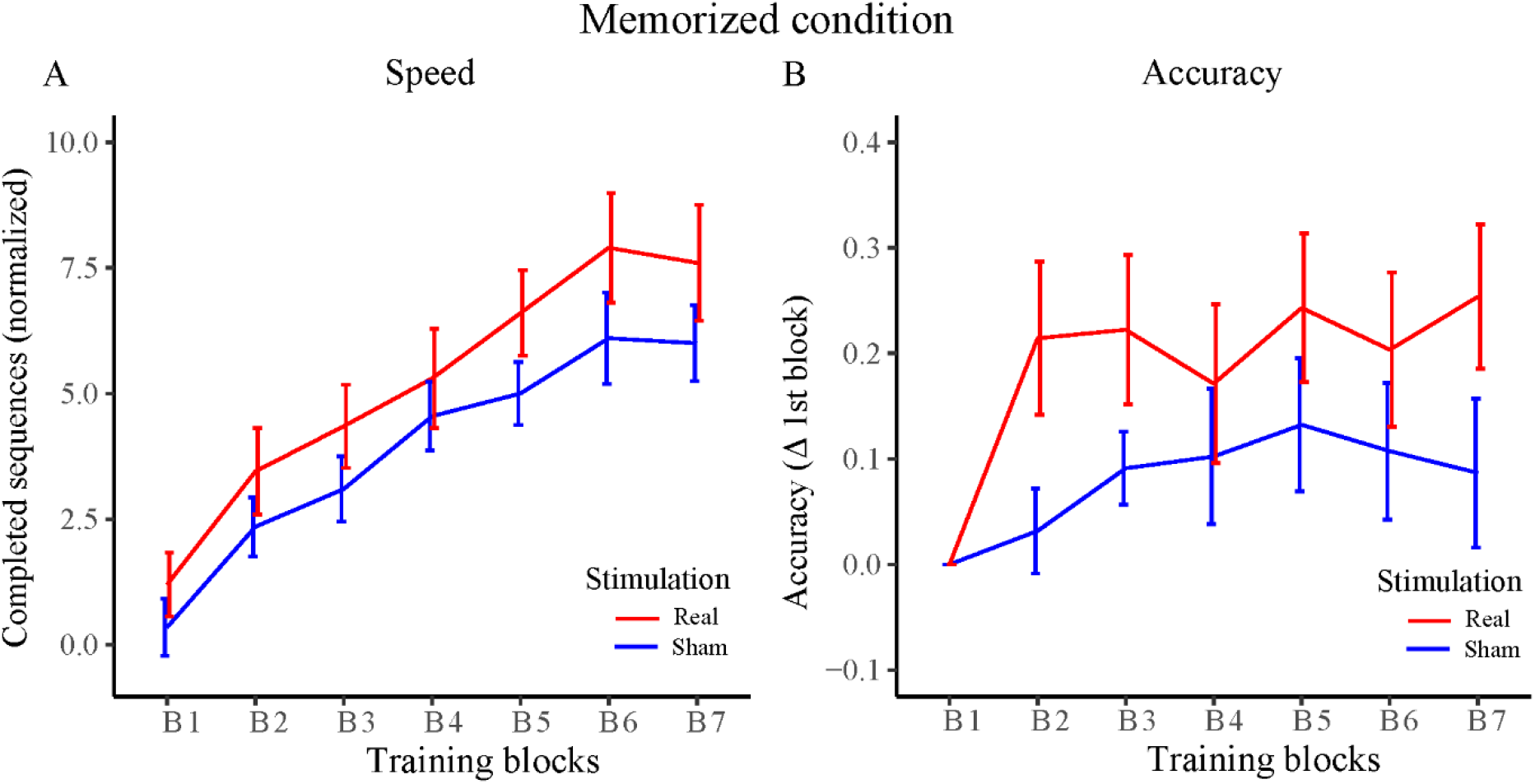
Plots of the speed and accuracy of the SFTT in the memorized condition. The error bars indicate standard error of the mean (SEM). Plot **A)** shows the speed determined by the total amount of completed sequences. Higher numbers depict better performance. Plot **B)** Shows the difference in accuracy as determined by the ratio of correct sequences divided by completed sequences, with regard to the first block. Please note the tACS significantly enhanced both, speed (A) and accuracy (B).

In order to see whether the increased amount of correct sequences was driven by an increase of speed or by a simultaneous increase of accuracy, we analysed the ratio between the total amount of sequences and the correct sequences as an accuracy measure. Upon visual inspection, the real stimulation group shows different dynamics in accuracy than the sham group. The real stimulation group demonstrates a steep increase in accuracy after the first training block while the sham group’s increase is more gradual. The accuracy between the groups during the 1^st^ training block was not significantly different T(19) = 0.85, *p* =.404. Therefore, to visualize the difference in dynamics we measured the difference in accuracy with regard to block 1. Results showed a significant block effect F(6, 247) = 3.47, *p* =.003, η_p_^2^ = 0.08 (medium effect), and a significant stimulation effect F(1, 247) = 18.31, *p* <.001, η_p_^2^ = 0.07 (medium effect), but no block x stimulation interaction F(6, 247) = 0.85, *p* = .529, η_p_^2^ = 0.02, see figure 4B. Indicating that speed as well as accuracy significantly improved with stimulation.

### N-back task

The N-back task performance was analysed by the following outcomes: hits, false alarms, accuracy (hits – false alarms), and reaction time for hits. All parameters were analysed separately using linear mixed effects models. The stimulation conditions (real vs. sham) and the three N-back difficulty levels were included as independent variables in the model. Two separate analyses were performed, one model included difficulty levels 1 and 2 to mimic the conditions comparable to the study of Violante *et al*. (2017), additionally we added difficulty level 3 to the model to test for a stimulation effect on the task with higher cognitive demand [27].

#### Reaction time

We were able to replicate the results of Violante and colleagues for the parameter reaction time with a significant effect of stimulation F(1, 47.10) = 5.33, *p* = .025, η_p_^2^ = 0.1 (medium effect), as well as an effect for difficulty level F(1, 35.77) = 44.61, *p* <.001 η_p_^2^ = 0.55 (large effect), and an interaction effect F(1, 34.54) = 4.83, *p* =.035, η_p_^2^ = 0.12 (medium effect). Post-hoc analysis with Tukey correction showed a significant difference between sham and real stimulation during difficulty level 2, t(42.1) = 3.22, *p* = .013, but not for level 1 t(42.6) = 0.20, *p* = .997. Both conditions showed a significant increase in reaction time between level 1 and level 2, which was more prominent in the sham condition t(36.4) = -6.16, *p* <.001 than the real condition t(36.4) = -3.25, *p* = .013 Adding the 3-back difficulty level to the model resulted in no effect for stimulation F(1,78.24) = 1.61, *p* = .209, a significant effect for difficulty level F(2, 67.65) = 34.99, *p* <.001 and no interaction effect F(2, 67.65) = 0.75, *p* = .478, see figure 5.

**Figure 5.**
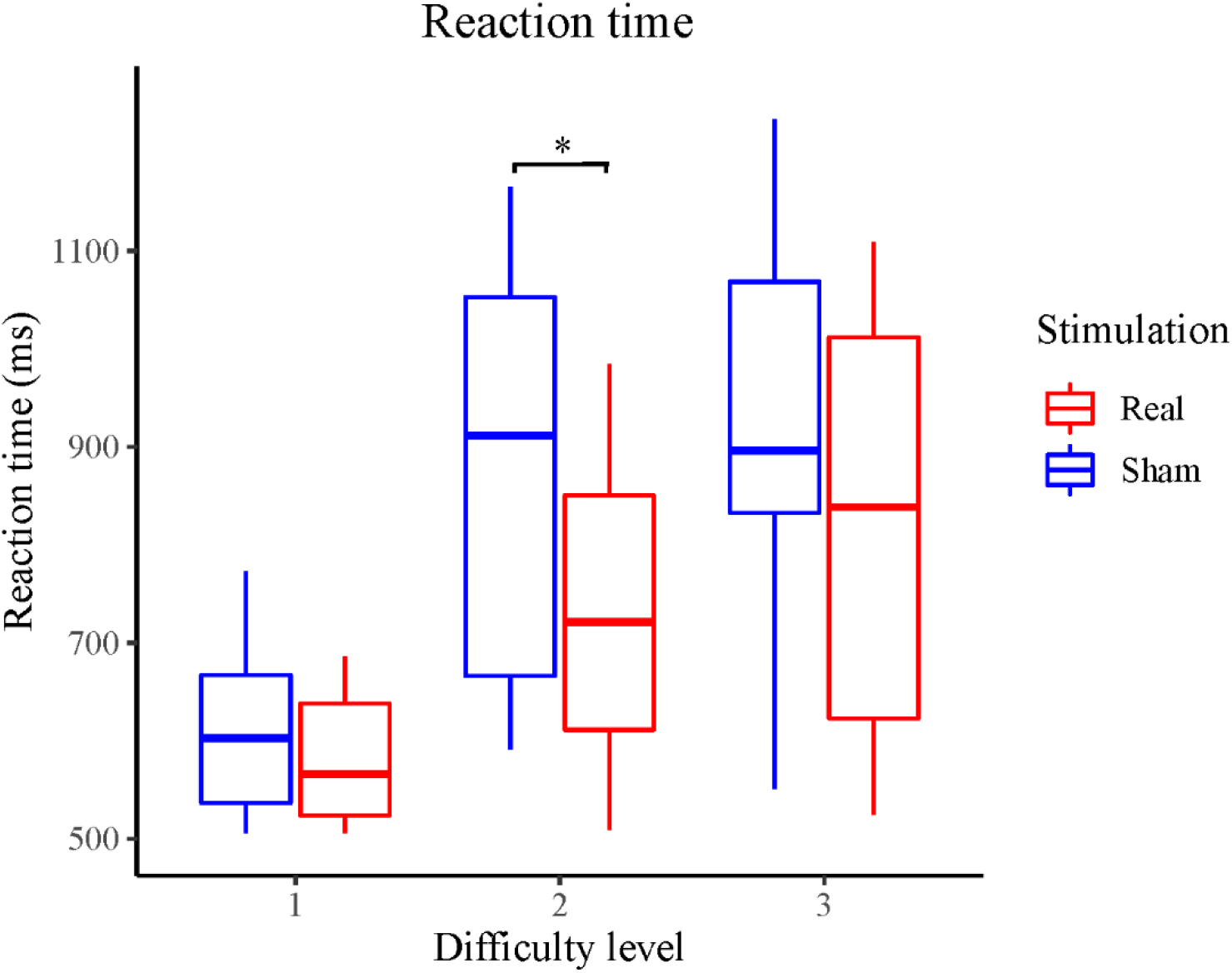
Figure of the reaction time for the hit responses on the N-back task. Difficulty levels 1-, 2- and 3-back included. The box ranges from Q1 (the 25^th^ percentile) to Q3 (the 75^th^ percentile), the bar shows the median. Results show better performance in reaction time during level 2-back for the real stimulation condition compared to sham.

#### N-back performance parameters (hits, false alarms, accuracy)

The analyses did not show a main effect of stimulation on any of these parameters for the 2 and the 3 level of difficulty models. There was a significant main effect of difficulty level. Indicating a significant decrease in performance with increasing n-back levels on all parameters. There were no stimulation x difficulty interaction effects. Please see table 1 for statistical results.

**Table 1.** N-back parameters.

| Parameter | Statistics |  |  |
| --- | --- | --- | --- |
| Model 1 | Stimulation | Difficulty | Interaction |
| Hits | $F(1, 40.4) = 0.80, p = .375$ | $F(1, 34.5) = 39.65, p < .001$ | $F(1, 33) = 0.49, p = .488$ |
| False alarms | $F(1, 33.1) = 0.54, p = .467$ | $F(1, 29.4) = 56.78, p < .001$ | $F(1, 29.4) = 0.87, p = .357$ |
| Accuracy | $F(1, 32.1) = 0.12, p = .733$ | $F(1, 25.9) = 85.12, p < .001$ | $F(1, 25.9) = 0.05, p = .823$ |
| Model 2 | Stimulation | Difficulty | Interaction |
| Hits | $F(1, 75.4) = 0.70, p = .407$ | $F(2, 65.2) = 69.46, p < .001$ | $F(2, 65.2) = 2.05, p = .137$ |
| False alarms | $F(1, 76) = 1.05, p = .309$ | $F(2, 65.8) = 42.26, p < .001$ | $F(2, 65.8) = 0.43, p = .651$ |
| Accuracy | $F(1, 75.5) = 0.02, p = .892$ | $F(2, 65.4) = 65.38, p < .001$ | $F(2, 65.4) = 1.57, p = .215$ |

Statistical results of all the n-back parameters. Columns show the main effect and the interaction effect of the independent variables. Model 1 shows the analysis with difficulty level 1 and 2, model 2 additionally includes level 3.

### Peak frequency analysis

During the EEG measurements data from 9 electrodes were acquired during the performance of the pre-baseline measurement of the N-back task. The individual average peak frequency of the 3 N-back levels combined was used as the personalized theta stimulation frequency for the rest of the study. Results of the overall average peak frequency showed a group mean of 4.5 (sd = 0.28) with a range between 4.1 and 5.4. For more details, please see table 2.

**Table 2.** Peak frequency analysis.

|  | Mean Theta | 1-back | 2-back | 3-back |
| --- | --- | --- | --- | --- |
| Mean (sd) | 4.48 (0.28) | 4.42 (0.35) | 4.46 (0.46) | 4.51 (0.65) |
| Minimum | 4.1 | 4 | 4 | 4 |
| Maximum | 5.4 | 5.7 | 6 | 7.8 |
| Missing | 0 | 0 | 0 | 1 |

Descriptive statistics of the peak frequency analysis measured during the three levels of the pre-baseline N-back measurements. The results show the overall means from the three levels combined and the means per N-back level. For the personalized stimulation paradigm, the individual mean peak frequency of the 3 N-back levels combined was used.

### Stimulation sensations & blinding

At the end of the last stimulation session, we investigated whether the stimulation was well tolerated and if there was a significant difference in experienced sensations between the real and sham condition. Moreover, we asked the participants to indicate whether they thought they had received real or sham stimulation during the before and after cross-over sessions.

The stimulation sensations were described with the use of a structured interview [59]. We checked for the following sensations: itching, pain, burning, metallic/iron taste in mouth, warmth, fatigue, other. With the possibility to respond: “none”, “mild”, “moderate”, “strong”. There were no adverse effects due to the tACS stimulation and only minor tACS sensations were reported. Most participants responded either with “none” or “mild”. Moreover, there was no significant difference between the stimulation and sham condition for any of the perceived sensations, see table 3.

**Table 3.** Stimulation sensations.

|  | None |  | Mild |  | Moderate |  | Strong |  | Statistics |  |
| --- | --- | --- | --- | --- | --- | --- | --- | --- | --- | --- |
|  | Real | Sham | Real | Sham | Real | Sham | Real | Sham | Chi-square | p-value |
| Itching | 75 | 80 | 20 | 15 | 5 | 5 | 0 | 0 | 0.18 | .916 |
| Pain | 85 | 85 | 5 | 15 | 10 | 0 | 0 | 0 | 3.00 | .223 |
| Burning | 80 | 80 | 10 | 20 | 10 | 0 | 0 | 0 | 2.67 | .264 |
| Warmth | 70 | 90 | 30 | 10 | 0 | 0 | 0 | 0 | 2.50 | .114 |
| Metallic/iron taste | 100 | 100 | 0 | 0 | 0 | 0 | 0 | 0 | n/a | n/a |
| Fatigue | 85 | 80 | 15 | 20 | 0 | 0 | 0 | 0 | 0.17 | .677 |
| Other | 80 | 80 | 15 | 15 | 5 | 5 | 0 | 0 | 0.00 | 1.000 |

*tACS sensations shown in percentages of participants who chose that specific response option for the intensity of the sensation. Statistics show comparison between real and sham stimulation.*

Participants were not able to discriminate between real and sham stimulation. A binomial test indicated that the proportion of correct answers during session 1 was 0.4, which was not significantly different than the chance level (0.5), *p* = .503. For session 2, the proportion of correct answers was 0.6, which was not significantly different than chance, *p* = .503, see table 4.

**Table 4.** Descriptive statistics for the tACS blinding.

| Session | Answer | Frequency | % |
| --- | --- | --- | --- |
| Before cross-over | Correct | 8 | 40 |
|  | Incorrect | 12 | 60 |
|  | Total | 20 | 100 |
| After cross-over | Correct | 12 | 60 |
|  | Incorrect | 8 | 40 |
|  | Total | 20 | 100 |

*The frequencies of correct and incorrect distinctions between real and sham stimulation before and after cross-over.*

## Discussion

The main outcome of this study is that personalized and synchronized tACS to the right FPN can enhance performance on the SFTT with high WM load in healthy older participants. This stimulation paradigm did not affect the performance on SFTT with low WM load. These findings indicate that the efficacy of bifocal theta tACS applied synchronously to DLPFC and PPC is dependent on the underlying cognitive state during the task. This result is further supported by the findings that tACS also improved N-back task performance specific to a difficulty level that was demanding enough.

### Motor task

The present results support the view of a causal effect of synchronised bifocal theta frequency oscillations applied to the right FPN on MSL. Correlative evidence of the activation of the FPN during finger tapping tasks has been previously shown using neuroimaging [60,61]. The meta-analysis of Witt and colleagues (2008) has shown that visually or self-paced finger tapping tasks induce concordant activity in the right DLPFC and the right inferior parietal cortex [61]. However, to the best of our knowledge, this was the first time the FPN was used as a target for a tACS paradigm with the intend to improve MSL.

We were able to demonstrate an improvement in MSL with this approach, but exclusively for the SFTT condition with high WM load (memorized condition). Therefore, the efficacy of the present orchestrated stimulation paradigm on motor behaviour was dependent on the amount of WM load during the task. This is in line with the study of Violante and colleagues (2017) that showed that theta tACS to the right FPN improved performance on a WM task, but only for the task with higher WM load [27]. This could be explained by the fact that the FPN shows more coherence in the theta range during WM tasks with high WM load [62]. Another possible explanation is that the involvement of the FPN is related to a specific sub-process of WM. WM can be roughly divided into three sub-processes: encoding, maintenance and retrieval [63]. The non-memorized SFTT condition required the participants to learn the sequence while performing the movements, which falls under the encoding phase. During the memorized SFTT condition, the participants needed to maintain and retrieve the previously learned sequence while performing the movements. A recent meta-analysis has shown that during the transition from encoding to maintenance and retrieval stages, the involvement of the FPN progressively increases. Therefore, it can be well hypothesized that the memorized SFTT condition benefits more from the FPN as a target while the non-memorized SFTT condition profits more from stimulation of other brain regions. For instance, the acquisition phase relies heavily on the dorsal attention network (DAN), which predominantly includes the frontal eye fields and the intraparietal sulci [63,64]. Moreover, studies have shown a high involvement of the M1 during the early stages of learning, with a reduction of activity to baseline when a sequence becomes explicitly known [65,66].

The involvement of the frontal and parietal areas during MSL has been well established, however their precise functional role is less clear [65,67–69]. MSL can be divided into three different learning phases: stage 1 for acquisition, stage 2 for consolidation and stage 3 for retention. The early learning phase relies more heavily on cognitive processes such as WM, showing an activation in the prefrontal cortex and parietal areas [70–72]. In this study, the efficacy of targeting the FPN to enhance performance on the MSL task is most likely specific to WM load. Studies that focused on the WM processes found that the FPN is associated with the maintenance and manipulation of information when theta oscillations were in synchrony between the two brain areas [26,62]. This might explain why the performance on the motor sequence task only improved during the high WM load task, where the participants had to perform the sequence from memory. Moreover, both accuracy and speed improved significantly in the memorized condition due to the tACS stimulation. However, the real stimulation induced a sharp increase in accuracy, while the sham group improved more gradually. Similar results have been shown in a study comparing real vs. sham anodal transcranial direct current stimulation (atDCS) applied to the M1 on a SFTT. Different age groups were compared and older adults showed a sharp increase in accuracy in the real stimulation group and a gradual increase in the sham group [73]. They argue that the active M1 stimulation facilitated the encoding and storage of the sequence in memory. In the current study, the stimulation target was the FPN and was effective in the memorized condition when the sequences were already learned. This result could be driven by an enhanced capacity to maintain and retrieve the previously learned sequence, due to the synchronization of theta oscillations in the FPN [63].

Although this has been the first time that both the DLPFC and the PPC have been targeted with the use of bifocal stimulation during a sequence learning task, studies have targeted the two areas separately. Stimulation of the DLPFC with the use of transcranial direct current stimulation (tDCS) has found heterogenous results for motor learning tasks [74–76]. A recent study using high-definition tDCS (HD-tDCS) to the DLPFC before the practice of an explicit MSL task found a decrease in performance after both anodal and cathodal HD-tDCS [74]. Hashemirad and colleagues applied anodal tDCS to the DLPFC and the PPC separately during an MSL task. This stimulation paradigm did not enhance performance in the trained right hand or untrained left hand for both brain areas [76]. Another study compared anodal, cathodal and sham tDCS applied to the PPC before, during and after an MSL task. Although the results did not show any stimulation effect during the training, the cathodal stimulation did inhibit reaction times of learned sequences 30 minutes after the end of training [69], which implicates a possible role of the PPC during the offline-learning phase of MSL. As there are currently only few results about the effects of NIBS on the FPN during motor tasks, more studies need to be conducted to provide more conclusive results. Moreover, the abovementioned studies all used tDCS paradigms, which rely on a different mechanism of action than tACS, which was used in the current study [22,77]. Therefore, the differences in results might rely entirely on the difference in stimulation techniques. However, based on the heterogenous results of unifocal tDCS to the frontal or parietal areas during MSL and the improvement in performance after bifocal stimulation, one might suggest that targeting the FPN as a network is more effective than targeting single brain sites.

### Personalized tACS

This study has proven the efficacy of personalized tACS stimulation in the theta range on MSL and cognitive function. Individual peak frequencies were measured while participants performed the N-back task. The averaged peak frequency was used with the idea to best exogenously synchronize the applied oscillatory active with the cortical oscillations in their endogenous frequencies measured during the relevant underlying brain state. Individualizing stimulation paradigms is important due to the inter-individual variability in endogenous frequencies [78]. For example, the average stimulation frequency in our study was 4.5 Hz while the range of the individual frequencies was between 4 and 7.8 Hz. Therefore, selecting one frequency for all participants might introduce heterogeneity in the efficacy of the stimulation [79]. Adjusting stimulation frequencies specifically to the endogenous oscillations enhances the power of oscillations [80]. Moreover, the endogenous oscillations constrain the effects of tACS stimulation as tACS fails to exogenously induce a frequency shift of oscillations of a non-endogenous frequency [79]. Therefore, personalization of stimulation paradigms seems to be important to circumvent this constraint and possibly reduce heterogeneity in the outcomes of tACS studies. However, due to the current lack of comparative studies of the efficacy of tACS with personalized compared with standardized frequencies to enhance MSL, no conclusive statements can be made. More research is necessary to explore the efficacy of personalization.

### N-back task

The reason for the use of the N-back task was twofold. First and foremost, as a way to measure the individual theta frequency while performing a WM task. Second, it was used as an additional control experiment to verify that the stimulation was indeed directed to the FPN and modulates a key function processed by the FPN. The behavioural results of the WM task support the notion that theta tACS to the FPN enhances WM performance [26,27]. As there was no neuroimaging data to confirm that the FPN was indeed targeted, a behavioural difference in WM performance provides correlational evidence.

In this study, we could replicate the observations of Violante et al. showing that exogenous synchronization of cortical oscillations in the theta range improved WM performance when cognitive demands were moderately high (2-back level) [27]. We have extended the results with showing that this was only applicable to level 2-back and not the more difficult 3-back level. The efficacy of the stimulation paradigm seems to follow an inverted u-shape in relationship to the difficulty of the task. The present study cohort were healthy older adults. Although it is currently unclear whether young adults would still benefit from the oscillatory synchronization during the 3-back task, one could speculate that the inverted u-shape with the peak at the 2-back task is age related.

Studies that have compared performance on WM tasks between young and healthy older adults have shown age related reduction in performance especially in tasks with high cognitive demand [81,82]. In response to high WM load, older adults show a relative hypoactivation in fronto-parietal regions compared to young adults [81,83]. The “Compensation-Related Utilization of Neural Circuits Hypothesis” (CRUNCH) provides a framework for this phenomenon; age-related hyperactivations are seen during tasks with low WM load due to reduced neural efficiency, with hypoactivation for tasks with high WM load due to reduced neural capacity [84]. Showing that older adults use compensatory mechanisms already with low WM load tasks (1-back) and are therefore not able to recruit the necessary neural resources during high WM load tasks (3-back) [81,82]. Heinzel and colleagues hypothesized that the change in neuronal activity is due to a decrease in FPN coupling; they showed that fronto-parietal connectivity decreased in older adults during 2-back and even more during 3-back tasks [71,85]. This could indicate that the difference in efficacy of stimulation between the 2-back and the 3-back tasks is related to the degree of deficient coupling of the FPN within these tasks and that the interventional approach with tACS could only sufficiently compensate these mechanisms for the 2-back task, but not any more for the 3-back task. The lack of improvement during the 1-back condition could indicate that the natural compensatory mechanisms are not sensitive to the effects of this stimulation paradigm. This points towards a specific efficacy that is dependent on the brain state caused by the amount of WM load.

### Future steps

The present study was a proof-of-principle study with the aim to investigate the involvement of the FPN in motor sequence learning. This study has a few limitations, which are discussed by means of suggestions for future studies. Firstly, multiple training sessions and/or a follow up session will enhance our understanding about the consolidation and possible retention of behavioural improvement. Motor learning encompasses multiple processes such as online and offline learning. Online learning is the improvement during the training of the task; offline learning happens after training and is a vital part of the consolidation of learned behaviour [65,86–88]. Multiple sessions will allow to investigate whether improvement continues with multiple training sessions and whether it retains during longer periods. Secondly, future studies should additionally include a standardized frequency (e.g., 6 Hz) as used in many other tACS studies, to compare with a personalized frequency [25–27]. Comparing the standardized to a personalized stimulation paradigm could provide more conclusive results about the importance of personalization to endogenous oscillatory activity. Lastly, to further personalize the approach future studies should personalize the placement of electrodes to the individual brain based on simulations. In the current study the electrode placement was defined by standardized locations using an EEG cap with the 10/20 system. We have used concentric electrodes and each montage consisted of a small circular centre electrode surrounded by a larger return electrode. This set-up has shown to improve focality compared with other electrodes such as the 5 × 5 cm rectangular electrodes or ring electrode set-ups with the return electrode on a separate region [89]. For a simulation of the electric field distribution, please see figure 2B. This improved focality highlights the importance of precision of the electrode placement as the stimulation is most effective close to the centre of the electrodes [90]. The currently used technique based on the 10-20-electrode system has been widely used in NIBS studies [91]. However, this is a standardized electrode placement system based on anatomical landmarks that can vary across participants [92]. A recent study of Scrivener and Reader compared the locations of the electrode placements with the use of an EEG cap with MRI images of the same participants. They found that the electrode placements deviated from the actual cortical locations with the smallest SD of 4.35 mm in frontal areas and the largest SD of 6.25 mm in the occipital and parietal areas [93]. These deviations are unlikely to result in any behavioural differences due to the focality of the stimulation. However, it does show that there is room for improvement in terms of precise definition of target locations and consistency in electrode placement. A way to improve precision is by using neuronavigation techniques guided by structural neuroimaging or with the use of functional MRI to pinpoint the exact target locations for stimulation [91].

### Conclusion

In conclusion, in this study, we were able to show a causal relationship between stimulating the FPN and improvements on MSL. Moreover, we were able to show distinctive efficacy of FPN synchronization for motor tasks with low- and high WM load, resulting in significant enhancement of performance in the motor task with high WM load, but no stimulation effects on the motor task with low WM load. The mechanisms of action point towards an effect of the stimulation paradigm on an improved capacity to maintain and manipulate the sequences. The current knowledge about using tACS to target frontal and parietal areas to improve MSL is limited. However, these results indicate that targeting the FPN as a network using personalized bifocal oscillatory stimulation is a promising approach. In addition, the present study showed that theta tACS applied to the FPN improved WM performance. This reveals an important interplay between the motor and cognitive domain pointing to it as a promising target for interventional strategies based on NIBS. However, to do this successfully, it is critically important that such an approach might only be effective when then cognitive load of a respective task is significantly high as demonstrated here by the WM load.

Taken together, personalized orchestrated bifocal tACS applied to the FPN might be a promising strategy to enhance motor sequence learning in healthy older adults and potentially in neurological patients showing deficits in motor learning.

## Author contributions

LD: Conceptualization, Design of experiment, Methodology, Validation, Data acquisition, Formal analysis, Writing – original draft, Writing – review & editing, Visualization, Project administration. MM: Validation, Randomization, Data acquisition. MW: Conceptualization, Project administration, Writing – review & editing. TM: Writing – Review & editing. FH: Conceptualization, Design of experiment, Interpretation of results, Writing – Review & editing.

## Funding

The present project was supported by the Defitech Foundation (Morges, CH).

## Acknowledgements

We thank Pablo Maceira for reading the manuscript and providing excellent comments, Elena Beanato for providing the script to pre-process the SFTT data, Roberto Salamanca-Giron for his contribution by providing and adapting his Matlab script for the EEG peak frequency analysis and Giorgia Giulia Evangelista for her contribution to the set-up of SimNIBS and her help with the visualization of the ring electrodes and the electric field distribution.

This study was supported by the EEG facility of the Human Neuroscience Platform, Fondation Campus Biotech Geneva, Geneva, Switzerland.

## Notes

### Competing Interest Statement

The authors have declared no competing interest.

## References

[1] Willingham DB. A neuropsychological theory of motor skill learning. Psychol Rev 1998;105:558–84. https://doi.org/10.1037/0033-295X.105.3.558.

[2] Dupont-Hadwen J, Bestmann S, Stagg CJ. Motor training modulates intracortical inhibitory dynamics in motor cortex during movement preparation. Brain Stimulat 2019;12:300–8. https://doi.org/10.1016/j.brs.2018.11.002.

[3] Karni A, Meyer G, Jezzard P, Adams MM, Turner R, Ungerleider LG. Functional MRI evidence for adult motor cortex plasticity during motor skill learning. Nature 1995;377:155–8. https://doi.org/10.1038/377155a0.

[4] Seidler RD, Bo J, Anguera JA. Neurocognitive contributions to motor skill learning: the role of working memory. J Mot Behav 2012;44:445–53. https://doi.org/10.1080/00222895.2012.672348.

[5] Buch ER, Santarnecchi E, Antal A, Born J, Celnik PA, Classen J, et al. Effects of tDCS on motor learning and memory formation: A consensus and critical position paper. Clin Neurophysiol Off J Int Fed Clin Neurophysiol 2017;128:589–603. https://doi.org/10.1016/j.clinph.2017.01.004.

[6] Wessel MJ, Zimerman M, Hummel FC. Non-invasive brain stimulation: an interventional tool for enhancing behavioral training after stroke. Front Hum Neurosci 2015;9:265. https://doi.org/10.3389/fnhum.2015.00265.

[7] Krause V, Meier A, Dinkelbach L, Pollok B. Beta Band Transcranial Alternating (tACS) and Direct Current Stimulation (tDCS) Applied After Initial Learning Facilitate Retrieval of a Motor Sequence. Front Behav Neurosci 2016;10. https://doi.org/10.3389/fnbeh.2016.00004.

[8] Pollok B, Boysen A-C, Krause V. The effect of transcranial alternating current stimulation (tACS) at alpha and beta frequency on motor learning. Behav Brain Res 2015;293:234–40. https://doi.org/10.1016/j.bbr.2015.07.049.

[9] Anguera JA, Reuter-Lorenz PA, Willingham DT, Seidler RD. Contributions of spatial working memory to visuomotor learning. J Cogn Neurosci 2010;22:1917–30. https://doi.org/10.1162/jocn.2009.21351.

[10] Maxwell JP, Masters RSW, Eves FF. The role of working memory in motor learning and performance. Conscious Cogn 2003;12:376–402. https://doi.org/10.1016/s1053-8100(03)00005-9.

[11] Krakauer JW, Hadjiosif AM, Xu J, Wong AL, Haith AM. Motor Learning. Compr Physiol 2019;9:613–63. https://doi.org/10.1002/cphy.c170043.

[12] Bo J, Borza V, Seidler RD. Age-Related Declines in Visuospatial Working Memory Correlate With Deficits in Explicit Motor Sequence Learning. J Neurophysiol 2009;102:2744–54. https://doi.org/10.1152/jn.00393.2009.

[13] Bo J, Seidler RD. Visuospatial Working Memory Capacity Predicts the Organization of Acquired Explicit Motor Sequences. J Neurophysiol 2009;101:3116–25. https://doi.org/10.1152/jn.00006.2009.

[14] Shea CH, Park J-H, Braden HW. Age-related effects in sequential motor learning. Phys Ther 2006;86:478–88.

[15] Baddeley AD, Hitch G. Working Memory. Psychol. Learn. Motiv., vol. 8, Elsevier; 1974, p. 47–89. https://doi.org/10.1016/S0079-7421(08)60452-1.

[16] Hikosaka O, Nakamura K, Sakai K, Nakahara H. Central mechanisms of motor skill learning. Curr Opin Neurobiol 2002;12:217–22.

[17] Pascual-Leone A, Wassermann EM, Grafman J, Hallett M. The role of the dorsolateral prefrontal cortex in implicit procedural learning. Exp Brain Res 1996;107:479–85.

[18] Verwey WB. Concatenating familiar movement sequences: the versatile cognitive processor. Acta Psychol (Amst) 2001;106:69–95. https://doi.org/10.1016/S0001-6918(00)00027-5.

[19] Bo J, Borza V, Seidler RD. Age-Related Declines in Visuospatial Working Memory Correlate With Deficits in Explicit Motor Sequence Learning. J Neurophysiol 2009;102:2744–54. https://doi.org/10.1152/jn.00393.2009.

[20] Shea CH, Park J-H, Braden HW. Age-related effects in sequential motor learning. Phys Ther 2006;86:478–88.

[21] Verhaeghen P, Cerella J. Aging, executive control, and attention: a review of meta-analyses. Neurosci Biobehav Rev 2002;26:849–57. https://doi.org/10.1016/S0149-7634(02)00071-4.

[22] Antal A, Paulus W. Transcranial alternating current stimulation (tACS). Front Hum Neurosci 2013;7:317. https://doi.org/10.3389/fnhum.2013.00317.

[23] Draaisma LR, Wessel MJ, Hummel FC. Non-invasive brain stimulation to enhance cognitive rehabilitation after stroke. Neurosci Lett 2020;719:133678. https://doi.org/10.1016/j.neulet.2018.06.047.

[24] Herrmann CS, Rach S, Neuling T, Struber D. Transcranial alternating current stimulation: a review of the underlying mechanisms and modulation of cognitive processes. Front Hum Neurosci 2013;7:279. https://doi.org/10.3389/fnhum.2013.00279.

[25] Kuo MF, Nitsche MA. Effects of transcranial electrical stimulation on cognition. Clin EEG Neurosci 2012;43:192–9. https://doi.org/10.1177/1550059412444975.

[26] Polania R, Paulus W, Nitsche MA. Modulating cortico-striatal and thalamo-cortical functional connectivity with transcranial direct current stimulation. Hum Brain Mapp 2012;33:2499–508. https://doi.org/10.1002/hbm.21380.

[27] Violante IR, Li LM, Carmichael DW, Lorenz R, Leech R, Hampshire A, et al. Externally induced frontoparietal synchronization modulates network dynamics and enhances working memory performance. ELife 2017;6:e22001. https://doi.org/10.7554/eLife.22001.

[28] Floyer-Lea A, Matthews PM. Distinguishable brain activation networks for short-and long-term motor skill learning. J Neurophysiol 2005;94:512–8. https://doi.org/00717.2004 [pii] 10.1152/jn.00717.2004.

[29] Honda M, Shibasaki H. Cortical control of complex sequential movement studied by functional neuroimaging techniques. Neuropathology 1998;18:357–62. https://doi.org/10.1111/j.1440-1789.1998.tb00131.x.

[30] Lin C-HJ, Chiang M-C, Wu AD, Iacoboni M, Udompholkul P, Yazdanshenas O, et al. Enhanced Motor Learning in Older Adults Is Accompanied by Increased Bilateral Frontal and Fronto-Parietal Connectivity. Brain Connect 2012;2:56–68. https://doi.org/10.1089/brain.2011.0059.

[31] Pammi VSC, Miyapuram KP, Ahmed, Samejima K, Bapi RS, Doya K. Changing the structure of complex visuo-motor sequences selectively activates the fronto-parietal network. NeuroImage 2012;59:1180–9. https://doi.org/10.1016/j.neuroimage.2011.08.006.

[32] Varela F, Lachaux JP, Rodriguez E, Martinerie J. The brainweb: phase synchronization and large-scale integration. Nat Rev Neurosci 2001;2:229–39. https://doi.org/10.1038/3506755035067550 [pii].

[33] Fries P. A mechanism for cognitive dynamics: neuronal communication through neuronal coherence. Trends Cogn Sci 2005;9:474–80. https://doi.org/10.1016/j.tics.2005.08.011.

[34] Fries P. Rhythms for Cognition: Communication through Coherence. Neuron 2015;88:220–35. https://doi.org/10.1016/j.neuron.2015.09.034.

[35] Fröhlich F. Noninvasive Brain Stimulation. Netw. Neurosci., Elsevier; 2016, p. 197–210. https://doi.org/10.1016/B978-0-12-801560-5.00015-X.

[36] Antal A, Herrmann CS. Transcranial Alternating Current and Random Noise Stimulation: Possible Mechanisms. Neural Plast 2016;2016:3616807. https://doi.org/10.1155/2016/3616807.

[37] Fröhlich F. Noninvasive Brain Stimulation. Netw. Neurosci., Elsevier; 2016, p. 197–210. https://doi.org/10.1016/B978-0-12-801560-5.00015-X.

[38] Constantinidis C, Klingberg T. The neuroscience of working memory capacity and training. Nat Rev Neurosci 2016;17:438–49. https://doi.org/10.1038/nrn.2016.43.

[39] Fell J, Axmacher N. The role of phase synchronization in memory processes. Nat Rev Neurosci 2011;12:105–18. https://doi.org/10.1038/nrn2979.

[40] Sauseng P, Klimesch W. What does phase information of oscillatory brain activity tell us about cognitive processes? Neurosci Biobehav Rev 2008;32:1001–13. https://doi.org/10.1016/j.neubiorev.2008.03.014.

[41] Zimerman M, Wessel MJ, Timmermann JE, Granstrom S, Gerloff C, Mautner VF, et al. Impairment of Procedural Learning and Motor Intracortical Inhibition in Neurofibromatosis Type 1 Patients. EBioMedicine 2015;2:1430–7. https://doi.org/10.1016/j.ebiom.2015.08.036.

[42] Haith AM, Krakauer JW. The multiple effects of practice: skill, habit and reduced cognitive load. Curr Opin Behav Sci 2018;20:196–201. https://doi.org/10.1016/j.cobeha.2018.01.015.

[43] Oldfield RC. The assessment and analysis of handedness: the Edinburgh inventory. Neuropsychologia 1971;9:97–113.

[44] World Medical Association. World Medical Association Declaration of Helsinki: ethical principles for medical research involving human subjects. JAMA 2013;310:2191–4. https://doi.org/10.1001/jama.2013.281053.

[45] Radloff LS. The CES-D Scale: A Self-Report Depression Scale for Research in the General Population. Appl Psychol Meas 1977;1:385–401. https://doi.org/10.1177/014662167700100306.

[46] Jaeggi SM, Studer-Luethi B, Buschkuehl M, Su Y-F, Jonides J, Perrig WJ. The relationship between n-back performance and matrix reasoning — implications for training and transfer. Intelligence 2010;38:625–35. https://doi.org/10.1016/j.intell.2010.09.001.

[47] Quent JA. JAQuent/nBack: Version 1.8. Zenodo; 2021. https://doi.org/10.5281/ZENODO.5502474.

[48] Vanderplas JM, Garvin EA. The association value of random shapes. J Exp Psychol 1959;57:147–54. https://doi.org/10.1037/h0048723.

[49] Wessel MJ, Park C, Beanato E, Cuttaz EA, Timmermann JE, Schulz R, et al. Multifocal stimulation of the cerebro-cerebellar loop during the acquisition of a novel motor skill. Sci Rep 2021;11:1756. https://doi.org/10.1038/s41598-021-81154-2.

[50] Lempel A, Ziv J. On the complexity of finite sequences. IEEE Trans Inf Theory 1976;22:75–81.

[51] Zimerman M, Heise KF, Gerloff C, Cohen LG, Hummel FC. Disrupting the ipsilateral motor cortex interferes with training of a complex motor task in older adults. Cereb Cortex 2014;24:1030–6. https://doi.org/10.1093/cercor/bhs385.

[52] Gandiga PC, Hummel FC, Cohen LG. Transcranial DC stimulation (tDCS): a tool for double-blind sham-controlled clinical studies in brain stimulation. Clin Neurophysiol 2006;117:845–50. https://doi.org/10.1016/j.clinph.2005.12.003.

[53] Thielscher A, Antunes A, Saturnino GB. Field modeling for transcranial magnetic stimulation: A useful tool to understand the physiological effects of TMS? 2015 37th Annu. Int. Conf. IEEE Eng. Med. Biol. Soc. EMBC, Milan: IEEE; 2015, p. 222–5. https://doi.org/10.1109/EMBC.2015.7318340.

[54] Salamanca-Giron RF, Raffin E, Zandvliet SB, Seeber M, Michel CM, Sauseng P, et al. Bifocal tACS Enhances Visual Motion Discrimination by Modulating Phase Amplitude Coupling Between V1 and V5 Regions. Neuroscience; 2020. https://doi.org/10.1101/2020.11.16.382267.

[55] Gravetter FJ, Wallnau LB. Essentials of statistics for the behavioral sciences. 8th Edition. Australia: Wadsworth, Cengage Learning; 2014.

[56] RStudio Team. RStudio: Integrated Development Environment for R. Boston, MA: RStudio, PBC; 2021.

[57] Kuznetsova A, Brockhoff PB, Christensen RHB. lmerTest Package: Tests in Linear Mixed Effects Models. J Stat Softw 2017;82. https://doi.org/10.18637/jss.v082.i13.

[58] JASP Team. JASP (Version 0.8.5.1). 2019.

[59] Antal A, Alekseichuk I, Bikson M, Brockmöller J, Brunoni AR, Chen R, et al. Low intensity transcranial electric stimulation: Safety, ethical, legal regulatory and application guidelines. Clin Neurophysiol Off J Int Fed Clin Neurophysiol 2017;128:1774–809. https://doi.org/10.1016/j.clinph.2017.06.001.

[60] Maruyama S, Fukunaga M, Sugawara SK, Hamano YH, Yamamoto T, Sadato N. Cognitive control affects motor learning through local variations in GABA within the primary motor cortex. Sci Rep 2021;11:18566. https://doi.org/10.1038/s41598-021-97974-1.

[61] Witt ST, Laird AR, Meyerand ME. Functional neuroimaging correlates of finger-tapping task variations: An ALE meta-analysis. NeuroImage 2008;42:343–56. https://doi.org/10.1016/j.neuroimage.2008.04.025.

[62] Sauseng P, Klimesch W, Schabus M, Doppelmayr M. Fronto-parietal EEG coherence in theta and upper alpha reflect central executive functions of working memory. Int J Psychophysiol 2005;57:97–103. https://doi.org/10.1016/j.ijpsycho.2005.03.018.

[63] Kim H. Neural activity during working memory encoding, maintenance, and retrieval: A network-based model and meta-analysis. Hum Brain Mapp 2019;40:4912–33. https://doi.org/10.1002/hbm.24747.

[64] Chabran E, Noblet V, Loureiro de Sousa P, Demuynck C, Philippi N, Mutter C, et al. Changes in gray matter volume and functional connectivity in dementia with Lewy bodies compared to Alzheimer’s disease and normal aging: implications for fluctuations. Alzheimers Res Ther 2020;12:9. https://doi.org/10.1186/s13195-019-0575-z.

[65] Dayan E, Cohen LG. Neuroplasticity subserving motor skill learning. Neuron 2011;72:443–54. https://doi.org/10.1016/j.neuron.2011.10.008.

[66] Pascual-Leone A, Grafman J, Hallett M. Modulation of cortical motor output maps during development of implicit and explicit knowledge [see comments]. Science 1994;263:1287–9.

[67] Doyon J, Gabitov E, Vahdat S, Lungu O, Boutin A. Current issues related to motor sequence learning in humans. Curr Opin Behav Sci 2018;20:89–97. https://doi.org/10.1016/j.cobeha.2017.11.012.

[68] Hardwick RM, Celnik PA. Cerebellar direct current stimulation enhances motor learning in older adults. Neurobiol Aging 2014;35:2217–21. https://doi.org/10.1016/j.neurobiolaging.2014.03.030.

[69] Pollok B, Keitel A, Foerster M, Moshiri G, Otto K, Krause V. The posterior parietal cortex mediates early offline-rather than online-motor sequence learning. Neuropsychologia 2020;146:107555. https://doi.org/10.1016/j.neuropsychologia.2020.107555.

[70] Anguera JA, Bernard JA, Jaeggi SM, Buschkuehl M, Benson BL, Jennett S, et al. The effects of working memory resource depletion and training on sensorimotor adaptation. Behav Brain Res 2012;228:107–15. https://doi.org/10.1016/j.bbr.2011.11.040.

[71] Heinzel S, Lorenz RC, Duong Q-L, Rapp MA, Deserno L. Prefrontal-parietal effective connectivity during working memory in older adults. Neurobiol Aging 2017;57:18–27. https://doi.org/10.1016/j.neurobiolaging.2017.05.005.

[72] Janacsek K, Nemeth D. Implicit sequence learning and working memory: Correlated or complicated? Cortex 2013;49:2001–6. https://doi.org/10.1016/j.cortex.2013.02.012.

[73] Maceira-Elvira P, Timmermann JE, Popa T, Schmid A-C, Krakauer JW, Morishita T, et al. Black-box testing in motor sequence learning. Neuroscience; 2021.https://doi.org/10.1101/2021.12.01.470563.

[74] Ballard HK, Eakin SM, Maldonado T, Bernard JA. Using high-definition transcranial direct current stimulation to investigate the role of the dorsolateral prefrontal cortex in explicit sequence learning. PLOS ONE 2021;16:e0246849. https://doi.org/10.1371/journal.pone.0246849.

[75] Gann MA, King BR, Dolfen N, Veldman MP, Davare M, Swinnen SP, et al. Prefrontal stimulation prior to motor sequence learning alters multivoxel patterns in the striatum and the hippocampus. Sci Rep 2021;11:20572. https://doi.org/10.1038/s41598-021-99926-1.

[76] Hashemirad F, Fitzgerald PB, Zoghi M, Jaberzadeh S. Single-Session Anodal tDCS with Small-Size Stimulating Electrodes Over Frontoparietal Superficial Sites Does Not Affect Motor Sequence Learning. Front Hum Neurosci 2017;11. https://doi.org/10.3389/fnhum.2017.00153.

[77] Nitsche MA, Paulus W. Excitability changes induced in the human motor cortex by weak transcranial direct current stimulation. J Physiol 2000;527 Pt 3:633–9.

[78] Fröhlich F, Riddle J. Conducting double-blind placebo-controlled clinical trials of transcranial alternating current stimulation (tACS). Transl Psychiatry 2021;11:284. https://doi.org/10.1038/s41398-021-01391-x.

[79] Schmidt SL, Iyengar AK, Foulser AA, Boyle MR, Fröhlich F. Endogenous Cortical Oscillations Constrain Neuromodulation by Weak Electric Fields. Brain Stimulat 2014;7:878–89. https://doi.org/10.1016/j.brs.2014.07.033.

[80] Boyle MR, Fröhlich F. EEG feedback-controlled transcranial alternating current stimulation. 2013 6th Int. IEEEEMBS Conf. Neural Eng. NER, San Diego, CA, USA: IEEE; 2013, p. 140–3. https://doi.org/10.1109/NER.2013.6695891.

[81] Nagel IE, Preuschhof C, Li S-C, Nyberg L, Bäckman L, Lindenberger U, et al. Load Modulation of BOLD Response and Connectivity Predicts Working Memory Performance in Younger and Older Adults. J Cogn Neurosci 2011;23:2030–45. https://doi.org/10.1162/jocn.2010.21560.

[82] Nyberg L, Dahlin E, Stigsdotter Neely A, Bäckman L. Neural correlates of variable working memory load across adult age and skill: Dissociative patterns within the fronto-parietal network. Scand J Psychol 2009;50:41–6. https://doi.org/10.1111/j.1467-9450.2008.00678.x.

[83] Rajah MN, D’Esposito M. Region-specific changes in prefrontal function with age: a review of PET and fMRI studies on working and episodic memory. Brain 2005;128:1964–83. https://doi.org/10.1093/brain/awh608.

[84] Reuter-Lorenz PA, Cappell KA. Neurocognitive Aging and the Compensation Hypothesis. Curr Dir Psychol Sci 2008;17:177–82. https://doi.org/10.1111/j.1467-8721.2008.00570.x.

[85] Heinzel S, Lorenz RC, Brockhaus W-R, Wustenberg T, Kathmann N, Heinz A, et al. Working Memory Load-Dependent Brain Response Predicts Behavioral Training Gains in Older Adults. J Neurosci 2014;34:1224–33. https://doi.org/10.1523/JNEUROSCI.2463-13.2014.

[86] Reis J, Schambra HM, Cohen LG, Buch ER, Fritsch B, Zarahn E, et al. Noninvasive cortical stimulation enhances motor skill acquisition over multiple days through an effect on consolidation. Proc Natl Acad Sci U A 2009;106:1590–5. https://doi.org/0805413106 [pii] 10.1073/pnas.0805413106.

[87] Robertson EM, Pascual-Leone A, Miall RC. Current concepts in procedural consolidation. Nat Rev Neurosci 2004;5:576–82. https://doi.org/10.1038/nrn1426.

[88] Robertson EM, Press DZ, Pascual-Leone A. Off-line learning and the primary motor cortex. J Neurosci 2005;25:6372–8. https://doi.org/10.1523/JNEUROSCI.1851-05.2005.

[89] Saturnino GB, Madsen KH, Siebner HR, Thielscher A. How to target inter-regional phase synchronization with dual-site Transcranial Alternating Current Stimulation. NeuroImage 2017;163:68–80. https://doi.org/10.1016/j.neuroimage.2017.09.024.

[90] Nitsche MA, Doemkes S, Karakose T, Antal A, Liebetanz D, Lang N, et al. Shaping the effects of transcranial direct current stimulation of the human motor cortex. J Neurophysiol 2007;97:3109–17. https://doi.org/01312.2006 [pii] 10.1152/jn.01312.2006.

[91] Woods AJ, Antal A, Bikson M, Boggio PS, Brunoni AR, Celnik P, et al. A technical guide to tDCS, and related non-invasive brain stimulation tools. Clin Neurophysiol 2016;127:1031–48. https://doi.org/10.1016/j.clinph.2015.11.012.

[92] Herwig U, Satrapi P, Schönfeldt-Lecuona C. Using the International 10-20 EEG System for Positioning of Transcranial Magnetic Stimulation. Brain Topogr 2003;16:95–9. https://doi.org/10.1023/B:BRAT.0000006333.93597.9d.

[93] Scrivener CL, Reader AT. Variability of EEG electrode positions and their underlying brain regions: visualising gel artifacts from a simultaneous EEG-fMRI dataset. Neuroscience; 2021. https://doi.org/10.1101/2021.03.08.434424.

